# Simple rules govern the diversity of bacterial nicotianamine-like metallophores

**DOI:** 10.1101/641969

**Authors:** Clémentine Laffont, Catherine Brutesco, Christine Hajjar, Gregorio Cullia, Roberto Fanelli, Laurent Ouerdane, Florine Cavelier, Pascal Arnoux

## Abstract

In metal-scarce environments, some pathogenic bacteria produce opine-type metallophores mainly to face the host’s nutritional immunity. This is the case of staphylopine, pseudopaline and yersinopine, identified in *Staphylococcus aureus*, *Pseudomonas aeruginosa* and *Yersinia pestis* respectively. These metallophores are synthesized by two (CntLM) or three enzymes (CntKLM), CntM catalyzing the last step of biosynthesis using diverse substrates (pyruvate or α-ketoglutarate), pathway intermediates (xNA or yNA) and cofactors (NADH or NADPH), depending on the species. Here, we explored substrate specificity of CntM by combining bioinformatics and structural analysis with chemical synthesis and enzymatic studies. We found that NAD(P)H selectivity was mainly due to the amino acid at position 33 (*S. aureus* numbering) which ensures a preferential binding to NADPH when it is an arginine. Moreover, whereas CntM from *P. aeruginosa* preferentially uses yNA over xNA, the staphylococcal enzyme is not stereospecific. Most importantly, selectivity towards α-ketoacids is largely governed by a single residue at position 150 of CntM (*S. aureus* numbering): an aspartate at this position ensures selectivity towards pyruvate whereas an alanine leads to the consumption of both pyruvate and α-ketoglutarate. Modifying this residue in *P. aeruginosa* led to a complete reversal of selectivity. Thus, opine-type metallophore diversity is mainly mediated by the absence/presence of a *cntK* gene encoding a histidine racemase, and the presence of an aspartate/alanine at position 150 of CntM. These two simple rules predict the production of a fourth metallophore by *Paenibacillus mucilaginosus*, which was confirmed *in vitro* and called bacillopaline.

## INTRODUCTION

In metal-scarce environments, bacteria have to use efficient mechanisms for the uptake of metals required for their growth. This is particularly the case for pathogenic bacteria that have to confront the host’s immune system. Indeed, the so-called “nutritional immunity” induces an additional metal limitation by sequestering iron, zinc or manganese to prevent bacterial growth [1–4]. To face this metal restriction, bacteria have developed metallophores to recover metals. In this context, nicotianamine-like metallophores have been identified in some bacteria as playing an important role in metal acquisition strategies. Staphylopine, pseudopaline and yersinopine are the three examples currently known and recently identified in *S. aureus* [5], *P. aeruginosa* [6,7] and *Y. pestis* [8] respectively. The biosynthesis of these nicotianamine-like metallophores occurs in two or three steps depending on the species. When it is present, as in *S. aureus*, CntK, a histidine racemase, transforms L-histidine (L-His) into D-histidine (D-His). CntL, a nicotianamine synthase-like, adds an aminobutyrate moiety coming from S-adenosyl methionine (SAM) on the amino group of the substrate (L-His in *P. aeruginosa* and *Y. pestis* or D-His in *S. aureus*) to form a pathway intermediate (yNA with L-His or xNA with D-His). Finally, CntM, an enzyme belonging to the opine dehydrogenase family, condenses the pathway intermediate with an α-ketoacid (pyruvate in the case of *S. aureus* and *Y. pestis* or α-ketoglutarate (αKG) in the case of *P. aeruginosa*) using NAD(P)H to form an opine-type metallophore (Figure 1).

**Figure 1:**
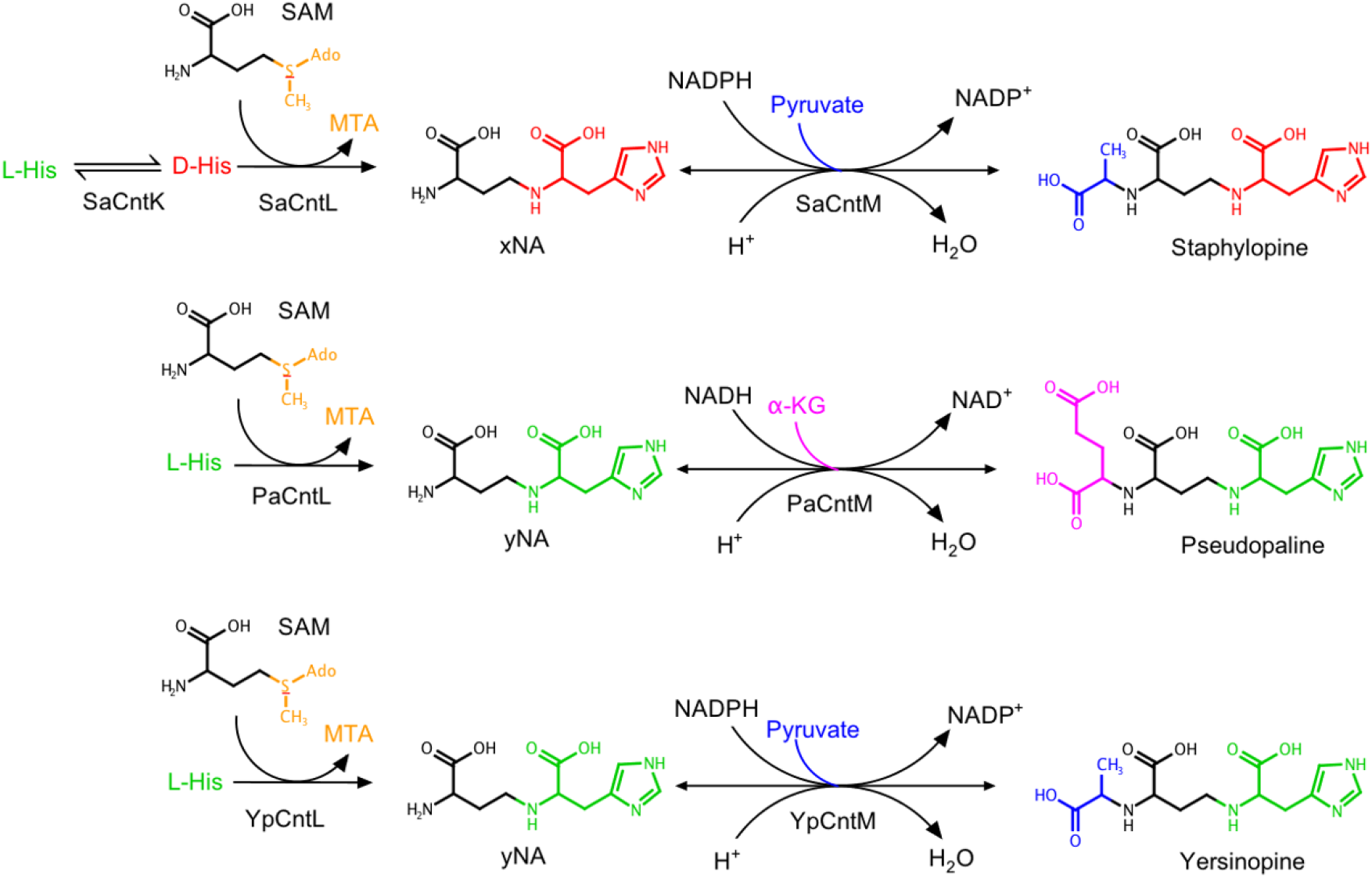
Differences in the staphylopine, pseudopaline, and yersinopine biosynthetic pathways. Adapted from [5,6,8].

Opine dehydrogenase catalyzes the NAD(P)H-dependent reductive condensation of the amino group of an amino acid with an α-ketoacid to produce an N-(carboxyalkyl) amino acid, also known as an opine, which exhibits either (L,L) or (D,L) stereochemistry [9]. The variety of amino acids and α-ketoacids that can be used as substrates by opine dehydrogenases results in diverse products. For example, the octopine dehydrogenase catalyzes the production of octopine, lysopine or histopine *via* reductive condensation of pyruvate and L-arginine, L-lysine and L-histidine respectively [10–13]. Leucinopine, asparaginopine and glutaminopine are other examples of opines produced *via* reductive condensation of α-ketoglutarate with L-leucine, L-asparagine and L-glutamine respectively [14,15]. In addition to the diversity of substrates, opine dehydrogenases are also distinguished by their biological roles. Indeed, in some marine invertebrates, opine dehydrogenases participate in the anaerobic metabolism by insuring the last step of anaerobic glycolysis pathway therefore participating in the propelling of these animals [16,17]. In plants, diverse opines are found inside crown gall tumors (for example nopaline, agropine, octopine, mannopine or D-L and L-L-succinamopine) that are induced by plant pathogenic bacteria as *Agrobacterium tumifaciens* [18]. In this case, the opines serve as nutrients conferring selective growth advantages to the opine-producing and opine-utilizing microorganisms [19,20].

Opine dehydrogenases are also involved in the biosynthesis of nicotianamine-like metallophores in bacteria and their substrate specificity results in the production of diverse metallophores. The activity of these opine dehydrogenases from *S. aureus*, *P. aeruginosa* and *Y. pestis* (respectively called SaCntM, PaCntM and YpCntM) has been described: SaCntM uses NADPH and pyruvate with xNA to produce staphylopine [5], YpCntM uses pyruvate and NADPH with yNA to produce yersinopine [8], and PaCntM uses NAD(P)H and α-ketoglutarate with yNA to produce pseudopaline [6,7]. Therefore, substrate specificity (and eventually stereospecificity in the case of xNA *vs* yNA) of CntM leads to the production of diverse opine-type metallophores. These metallophores are involved in metal acquisition, with metal specificity depending on the growth medium. Staphylopine could transport copper, nickel, cobalt, zinc and iron [5,21,22], and participates in zinc uptake in zinc-scarce environments [21,23]. Pseudopaline is involved in nickel uptake in minimal media, whereas it is responsible for zinc uptake in zinc-scarce environment [6,24]. For both *S. aureus* and *P. aeruginosa*, literature reports a link between the production of opine-type metallophores and bacterial infection. For example, in *S. aureus*, the staphylopine’receptor CntA plays an important role in the optimal functioning of the urease activity and in the virulence: deletion of this substrate binding protein leads to a decrease of murine bacteremia and urinary tract infections [22]. In *P. aeruginosa*, transcriptomic analyses showed that the biosynthetic genes for pseudopaline are overexpressed in burn wound infections in humans [25]. This overexpression would allow bypassing metal limitations set up during nutritional immunity. Moreover, the pseudopaline’s exporter CntI plays an important role in the survival and growth of *P. aeruginosa* in cystic fibrosis airway: deletion of this exporter results in an attenuation of this respiratory infection [26]. Similarly, the exporter of staphylopine was found to be important for fitness in abscesses, even before staphylopine discovery [27]. Concerning *Y. pestis*, no data indicates a link between yersinopine production and virulence at this time and the discovery of this metallophore is so far restricted to *in vitro* studies [8].

In an effort to understand bacterial opine-type metallophore diversity, we studied substrate specificity of CntM in *S. aureus* and *P. aeruginosa* by combining bioinformatic and structural analyses with chemical synthesis and enzymatic studies. A single amino acid residue is responsible for the preferential binding of NADPH, and a single residue is involved in the specificity towards pyruvate or α-ketoglutarate. These simple rules in substrate specificity prompted us to dig into available genomes for a bacteria possessing a *cntK* homologue together with a *cntM* gene predicted to use α-ketoglutarate. This research ultimately led to the discovery of a new opine-type metallophore called bacillopaline in *Paenibacillus mucilaginosus*.

## MATERIALS AND METHODS

### Bioinformatic analyses

SaCntM (*sav2468*) protein sequence was analyzed by searching for homologues using Psi-BLAST search [28] through the NCBI databases (National Center for Biotechnology Information, Bethseda, Maryland, USA) and the Pfam database (European Bioinformatics Institute, Hinxton, England, UK). Sequence alignment was done using the Muscle program [29] with defaults criteria from Jalview (version 2.10.3) [30]. Residues were colored following the Clustal coloring scheme with intensity modified by a conservation color increment of 30%. Gene syntheny were inspected using the MaGe MicroScope web interface [31] added to sequence alignment analysis.

### Cloning, expression and purification of proteins

The gene encoding the SaCntM protein (*sav2468*) was cloned in pET-SUMO and pET-101, and the one encoding the PaCntM protein (*pa4835*) was cloned in pET-TEV according to standard protocols. Similarly, the genes encoding the PmCntL and PmCntM proteins were cloned in pET-TEV from genomic DNA. The primers used for these constructions are listed in Table S1. After co-transformation with plasmid pRARE (encoding for rare codon in *E. coli*), *E. coli* BL21 strains were aerobically cultivated with horizontal shaking in LB media supplemented with appropriate antibiotics (kanamycin at 50 µg.mL^−1^ for pET-SUMO and pET-TEV, ampicillin at 50 µg.mL^−1^ for pET-101 and chloramphenicol at 25 µg.mL^−1^ for pRARE). These strains were grown in diverse conditions (37°C or 16°C, with or without induction of protein expression by addition of 0.1 mM IPTG when the OD of the culture was about 0.6; Table S2). After overnight growth, cells were recovered by centrifugation at 5,000 g for 20 min at 4°C. Cells were resuspended in buffer A and disrupted using a Constant cell disruption system operating at 1.9 Kbar. Cell debris were removed by centrifugation at 8,000 *g* for 20 min. The supernatant was centrifuged at 100,000 *g* for 45 min at 4°C to remove cell wall debris and membrane proteins. The resulting soluble fraction was purify by batch using a nickel-charged affinity resin (Ni-NTA Agarose resin, ThermoFisher Scientific). The proteins were eluted stepwise with imidazole (15 mM wash, 250 mM or 500 mM elution). Collected fractions were transferred into imidazole-free buffer B (see Table S2 for details on the buffers used).

### Site directed mutagenesis

Site directed mutagenesis were performed according to standard protocol from the QuickChange II Site-Directed Mutagenesis kit (Agilent Technologies). The only difference was that *E. coli* DH10β Competent Cells were used instead of XL1-Blue Supercompetent Cells for transformations. Selection of colonies was done after spreading on LB plate supplemented with appropriate antibiotics. Plasmid pET-SUMO containing the gene encoding the SaCntM protein and plasmid pET-TEV containing the gene encoding the PaCntM protein were used as mutagenesis templates for the D150A and A153D substitutions respectively. For the R33H substitution from SaCntM, mutagenesis was performed from plasmid pET-101 containing the gene encoding the SaCntM protein as template. Primer pairs were designed for single substitutions and were then synthesized by Eurofins Genomics (Table S1).

### Chemical synthesis of xNA and yNA

The chemical synthesis of xNA has recently been described [32]. The same strategy was used for the chemical synthesis of yNA, although L-His-OMe was used instead of D-His-OMe as starting material. Purified intermediates and their characterization are described in the supplementary materials.

### CntM activity assay

Enzymatic reactions of CntM were performed at 28°C in microplates and with purified proteins. They were carried out in a reaction volume of 100 µL in buffer C (50 mM BisTrisPropane, 100 mM NaCl, pH = 7 for SaCntM and PmCntM or pH = 8 for PaCntM) containing 2 or 5 µg of enzyme, 0.2 mM of NADH or NADPH (Sigma-Aldrich), 0.2 mM of xNA or yNA (chemically synthesized), and 1 mM of pyruvate or α-ketoglutarate (Sigma-Aldrich), unless otherwise stated. The absorbance at 340 nm was measured using a microplate reader (Infinite 200 Pro; Tecan) to follow the oxidation of NADPH illustrating the progress of the reaction. Activities were calculated from the initial rate and the amount of enzyme used. Kinetic parameters were estimated according to the Michaelis-Menten kinetic without **(1)** or with the substrate inhibition model **(2)** using SigmaPlot. These values were used to plot a fit on the experimental data. (*V*_*m*_: Maximum velocity; *K*_*m*_: Michaelis constant; [S]: substrate concentration; *K*_*i*_: Inhibition constant).

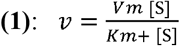

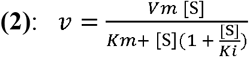

### Fluorescence resonance energy transfer (FRET) studies

Fluorescence studies were performed using a Cary Eclipse spectrophotometer (Agilent). The FRET experiment was done using a protein concentration of 5 μM and an excitation wavelength at 280 nm (tryptophan excitation). The emission at 340nm was transferred to the NAD(P)H and the signal was recorded between 400 and 500 nm. Five spectra were averaged in order to increase signal to noise ratio.

### Activity assay followed by TLC

The assay consisted in incubating the purified enzymes at a final concentration of 2.5 μM and using carboxyl-[^14^C]-labeled SAM (2.5 μM), NADPH or NADH (30 μM), L- or D-histidine (10 μM) and α-keto acid (pyruvate or α-ketoglutarate; 1mM). The total volume was 100 μL in buffer D (50 mM of Hepes, 1 mM of DTT, 1mM of EDTA, pH = 9). The mixtures were incubated for 30 min at 28 °C. The reactions were stopped by adding ethanol to a final concentration of 50 % (*v/v*) and the products were then separated by thin layer chromatography. An aliquot of 10 μL of the reaction mixtures were spotted on HPTLC (High Performance TLC) Silica Gel 60 Glass Plates (Merck KGaA), and the plates were developed with a phenol:*n*-butanol:formate:water (12:3:2:3 *v/v*) solvent system. These separation parameters were used in the initial biochemical characterization of plant nicotianamine synthase [33] and of staphylopine and pseudopaline [5,6]. HPTLC plates were dried and exposed to a [^14^C]-sensitive imaging plate for one day. Imaging plates were then scanned on a Typhoon FLA 7000 phosphorimager (GE Healthcare).

### Cell culture conditions of *Paenibacillus mucilaginosus*

*Paenibacillus mucilaginosus*, sub sp. 1480D (from Collection DSMZ) was grown by inoculating stock bacteria into 20 mL sterile medium (TSB 1/10) at 30°C for 48h. Genomic DNA was extracted using the protocols and pretreatments for Gram-positive bacteria from DNeasy Blood and Tissue kit (Qiagen).

## RESULTS AND DISCUSSION

### Specificity towards NADH or NADPH

The structure of CntM have been solved in a binary/tertiary complex with NADPH or NADPH and xNA, revealing the residues that are involved in the complex formation and building the active site [8,32]. First focusing on the NADPH binding site of SaCntM (Figure 2A) we observed that the phosphate group of NADPH is sandwiched by the side chains of two positively charged residues (R33 and K39) that are rather conserved in the CntM family. However, a sequence alignment of CntM from nine different species including *S. aureus*, *P. aeruginosa* and *Y. pestis* shows that R33 residue in SaCntM (also present in sequence from *Y. pestis*) is replaced by a histidine in PaCntM (Figure 2B). Because it is known that SaCntM and YpCntM specifically uses NADPH [5,8] while PaCntM could use NADH or NADPH depending on the conditions [6–8], we hypothesized that the nature of the amino acid residue at this position could determine the NADH/NADPH selectivity by CntM.

**Figure 2:**
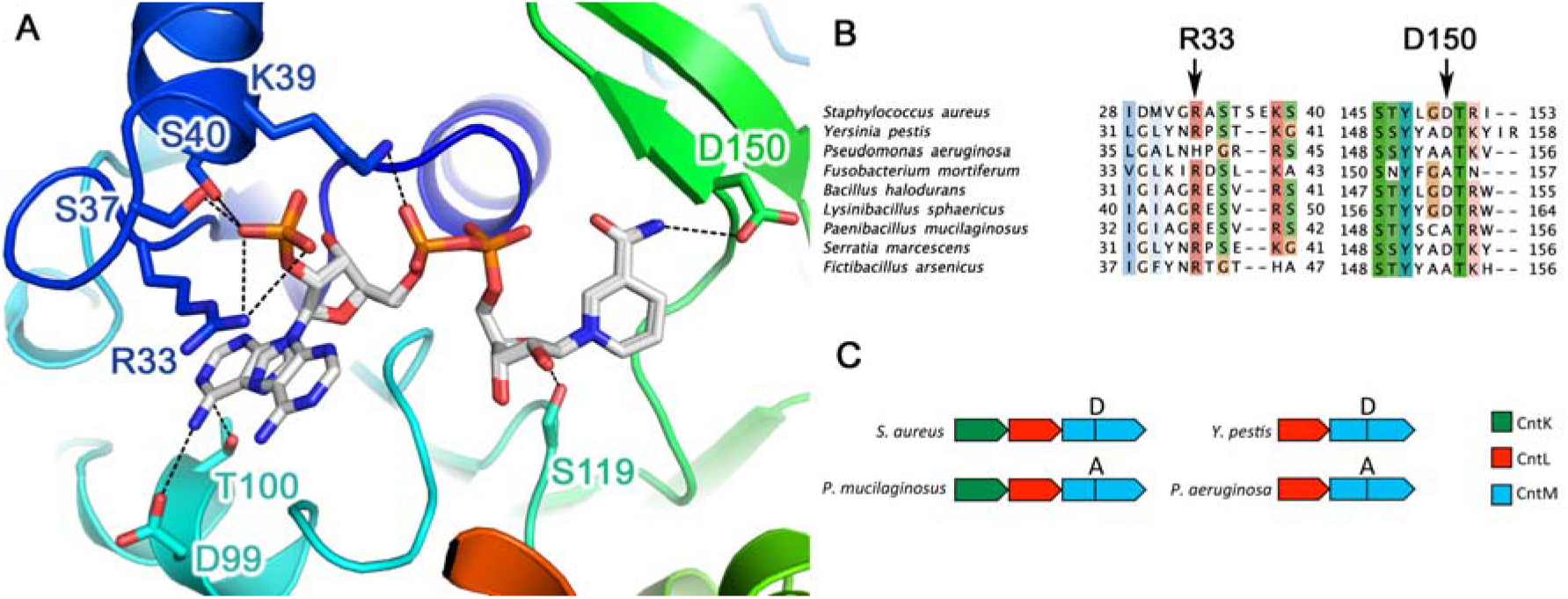
CntM structure, sequence conservation around NADPH and operon diversity highlight protein selectivity. (A) Details of the NADPH binding site on SaCntM with side chains of residues located up to 4Å around the NADPH represented in stick. The protein is colored from N-terminus (blue) to C-terminus (red). (B) Sequence alignment of nine CntM protein sequences from bacteria. Threshold for the Clustal coloring scheme correspond to 30 % sequence conservation as defined in Jalview. Arrows point to residues involved in NADPH/NADH selectivity and pyruvate/α-ketoglutarate selectivity. (C) Genomic organization of the biosynthetic genes of four different *cnt* operons in bacteria.

We therefore sought to determine the role of this residue in NAD(P)H selectivity by replacing this arginine by a histidine in a SaCntM:R33H variant. Because the histidine could either be neutral or positively charged at pH above or below its p*K*_a_, the binding of NAD(P)H was followed at two pHs (6.0 and 8.5). We found that the WT enzyme, whatever the pH, still preferentially binds NADPH over NADH (Figure 3). On the contrary, the R33H mutant behaves as the WT at pH 6.0, whereas it loses its preferential binding property at a higher pH. This shows that a histidine at this position could serve as a selective residue, stabilizing NADPH at acidic pH when the imidazole ring of histidine is positively charged, and favoring NADH binding when histidine is neutral at basic pH, overall explaining the difference of selectivity in the literature [6–8]. Interestingly, we noted that *Fictibacillus arsenicus* possesses a histidine residue at the conserved position equivalent to K39 in *S. aureus*, suggesting the same possibility of preferential NADH binding at basic pH.

**Figure 3:**
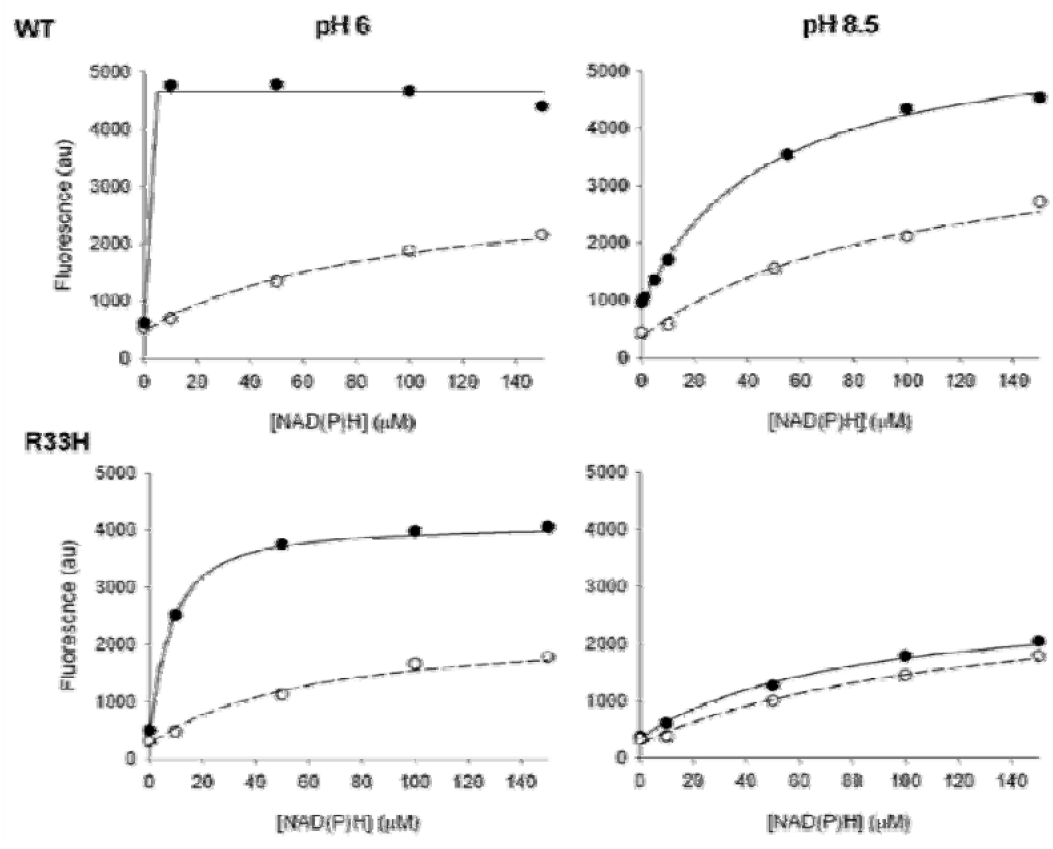
Titration of NADPH (black circles) and NADH (white circles) binding to SaCntM (5 µM) at pH = 6.0 or pH = 8.5 followed by fluorescence energy transfer between tryptophan excitation (280 nm) and NAD(P)H emission (450 nm).

### Specificity towards xNA or yNA

*In vivo*, CntM uses the product of CntL *i.e.* xNA or yNA depending on the species. However, using enzymatically produced xNA or yNA, McFarlane *et al.* (2018) suggested that SaCntM could use both diastereoisomers. Activities from *S. aureus* (SaCntM) and from *P. aeruginosa* (PaCntM) were therefore compared for their ability to use chemically synthesized xNA and yNA as substrate (Figure 4). The chemical synthesis of xNA was recently reported [32] and we were able to synthesize yNA by following the same approaches (see the experimental procedures and the supplementary materials). Reactions were then performed *in vitro* using purified proteins and a concentration range of xNA and yNA with a fixed concentration of other substrates: 0.2 mM of NADPH and 1 mM of pyruvate (when evaluating SaCntM) or α-ketoglutarate (when evaluating PaCntM). Overall, we found that the activity of CntM from *S. aureus* and *P. aeruginosa* towards yNA and xNA were different. In the case of SaCntM, although xNA is used *in vivo* to produce staphylopine, its activity is higher when using yNA *in vitro*. Indeed, the *k*_cat_ for the reaction with yNA is ~2-fold higher than the one with xNA (3.18 s^−1^ and 1.67 s^−1^ respectively). However, the *K*_m_ for the reaction with yNA is ~2-fold higher than the one with xNA (47 µM and 23 µM respectively), leading to a catalytic efficiency (*k*_cat_/*K*_m_) of the same order of magnitude for the two substrates (Table 1). In the case of PaCntM, the enzyme uses yNA to produce pseudopaline *in vivo*. In agreement with this, we found that the reaction with yNA is more efficient than the one with xNA, the reaction with yNA exhibiting a catalytic efficiency ~10-fold higher than with xNA (29 618 M^−1^s^−1^ and 2 918 M^−1^s^−1^ respectively).

**Table 1:** Kinetic parameters of SaCntM and PaCntM activities established for a concentration range of xNA and yNA with fixed concentrations of other substrates: 0.2 mM of NADPH and 1 mM of pyruvate when evaluating SaCntM, and 0.2 mM of NADH and 1 mM of α-ketoglutarate when evaluating PaCntM. The data and the standard errors associated with were generated by SigmaPlot according to the Michaelis-Menten model with or without substrate inhibition. (NA: Not Applicable; *V*_m_: Maximum velocity; *K*_m_: Michaelis constant; *K*_i_: Inhibition constant; *k*_cat_: Catalytic constant (or turnover number); *k*_cat_/*K*_m_: Catalytic efficiency).

| Protein | xNA vs yNA | $K_m$ ($\mu\text{M}$) | $K_i$ (mM) | $k_{\text{cat}}$ ($\text{s}^{-1}$) | $k_{\text{cat}}/K_m$ ($\text{M}^{-1}\text{s}^{-1}$) |
| --- | --- | --- | --- | --- | --- |
| SaCntM | yNA | $47 \pm 9$ | $2.3 \pm 0.8$ | $3.18 \pm 0.25$ | 67 584 |
| SaCntM | xNA | $23 \pm 3$ | NA | $1.67 \pm 0.04$ | 72 711 |
| PaCntM | yNA | $21 \pm 7$ | $2.4 \pm 1.0$ | $0.62 \pm 0.06$ | 29 618 |
| PaCntM | xNA | $54 \pm 20$ | NA | $0.16 \pm 0.02$ | 2 918 |

**Figure 4:**
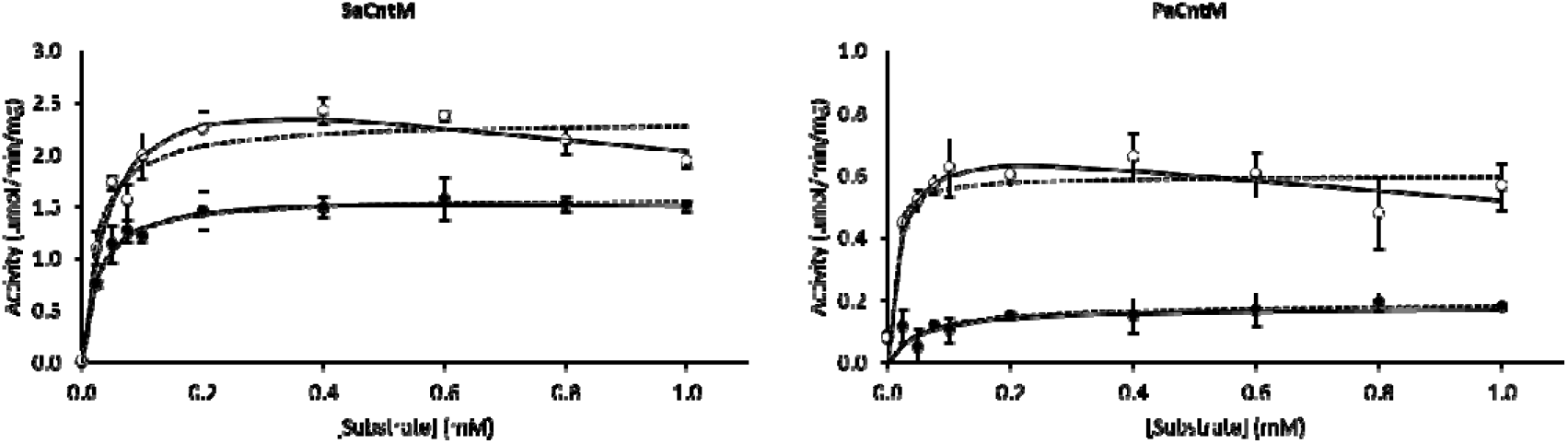
Activity profile of SaCntM and PaCntM using variable concentrations of xNA (black circle) and yNA (white circle) with fixed concentrations of other substrates: 0.2 mM of NADPH and 1 mM of pyruvate when evaluating SaCntM, or 0.2 mM of NADH and 1 mM of α-ketoglutarate when evaluating PaCntM. The data points are means of three replicates with standard deviations. The fits are made using the Michaelis-Menten model considering (continuous line) or not (dashed line) a substrate inhibition.

In the past decades, several opine dehydrogenases have been studied and their biosynthetic reactions generally exhibit a substrate stereospecificity towards the amino group with most enzymes using the L-stereoisomer as substrate [9]. For example, substrate stereospecificity has been outlined for the octopine dehydrogenase from *Pecten maximus* [12] and the structure of this enzyme shows a negatively charged cavity acting as a “charge ruler”, which favors L-arginine binding. Here, we found that even if SaCntM uses xNA *in vivo*, it is also capable of using yNA *in vitro*. This trend confirms the one outlined by McFarlane *et al.* [8] using enzymes from different species. Under their experimental conditions, PaCntM exhibited a *k*_cat_ of 0.016 s^−1^ in the presence of xNA, which was biosynthesized with SaCntL. Accordingly, we found that PaCntM preferentially used yNA whether *in vitro* or *in vivo.* All these data therefore suggest that there is a substrate stereospecificity in the case of PaCntM, which is not found in SaCntM. We further noted that, using yNA but not xNA, a drop in enzyme activity is visible at high substrate concentrations, which suggests a mechanism of substrate inhibition (Figure 4). Indeed, the fits made with a substrate inhibition model added to the Michaelis-Menten kinetic better cover the experimental data (Figure 4; plain lines). However, this substrate inhibition is not visible when using xNA as substrate.

### Specificity towards pyruvate or α-ketoglutarate

In order to find amino acid residues involved in the pyruvate/α-ketoglutarate selectivity by CntM, we searched for residues located in the vicinity of the active site (*i.e.* the nicotinamide moiety) and conserved in species known to use pyruvate (*S. aureus* and *Y. pestis*) but differing in species known to use α-ketoglutarate *(P. aeruginosa*). This pointed to the aspartic acid residue at position 150 in SaCntM, which is also present in YpCntM but replaced by an alanine in PaCntM (corresponding to residue 153 in this enzyme). We hypothesized that the nature of the amino acid residue at this position would determine the pyruvate/α-ketoglutarate selectivity by CntM. This postulate was also proposed by McFarlane *et al.* [8]. In order to test the role of this amino acid in pyruvate/α-ketoglutarate selectivity, we replaced the aspartic acid by an alanine in SaCntM (D150A variant) and did the opposite mutation in PaCntM (A153D). The activities of these proteins were then compared for their ability to use of pyruvate and α-ketoglutarate as substrates (Figure 5). The reactions were performed using a concentration range of pyruvate and α-ketoglutarate with a fixed concentration of others substrates: 0.2 mM of NAD(P)H and 0.2 mM of xNA (when using SaCntM) or yNA (PaCntM). As a control, we verified that the mutation did not affect the binding of the NADPH on SaCntM (Figure S8).

**Figure 5:**
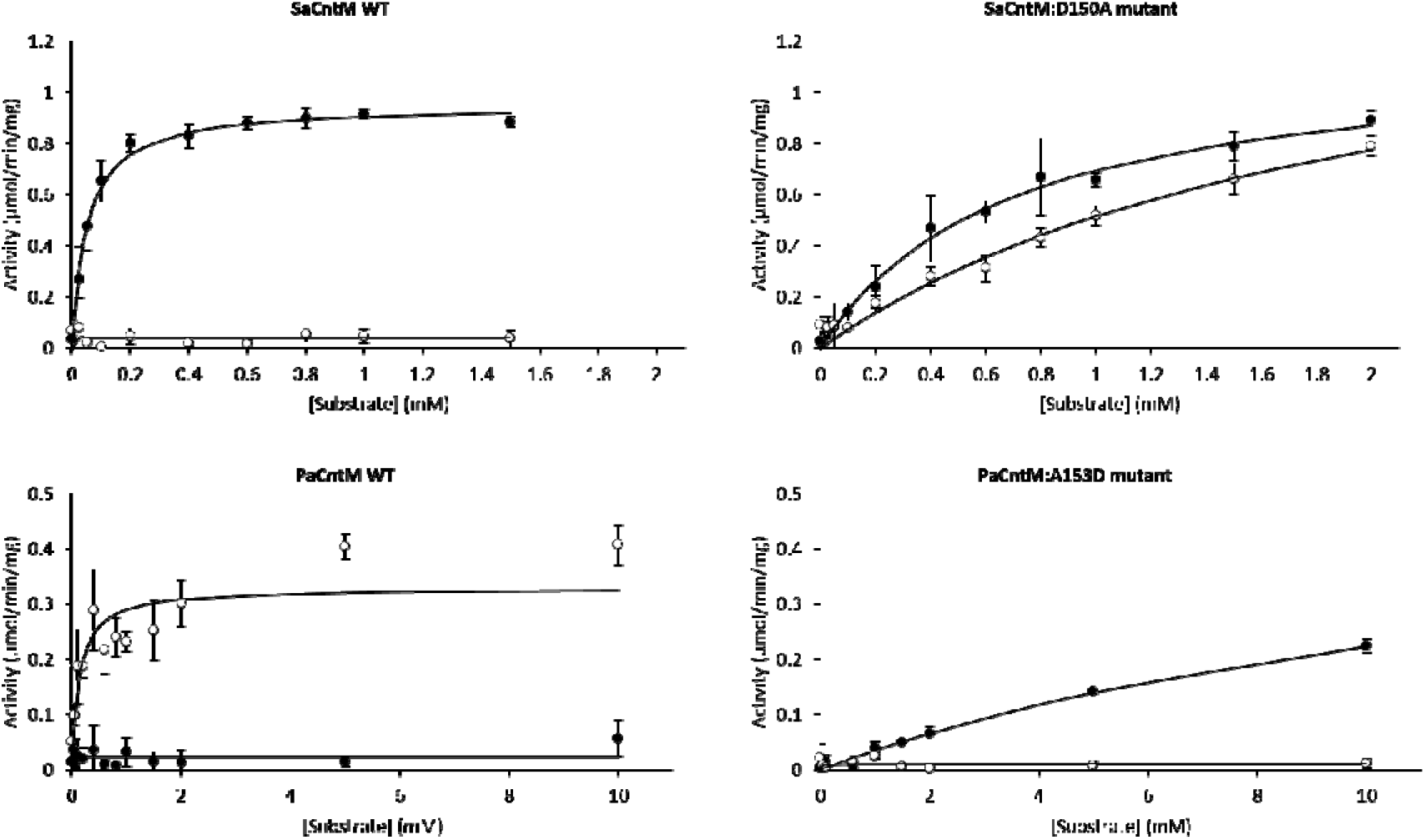
Activity profile of SaCntM (WT and D150A mutant) and PaCntM (WT and A153D mutant) using variable concentrations of pyruvate (black circle) and α-ketoglutarate (white circle) with fixed concentrations of others substrates: 0.2 mM of NADPH and xNA when evaluating SaCntM, and 0.2 mM of NADH and yNA when evaluating PaCntM. The data points are means of three replicates with standard deviations. The fits are made using the Michaelis-Menten model.

With regard to SaCntM, we confirmed that the WT was only able to use pyruvate, with a *k*_cat_ of 1.0 s^−1^ and a *K*_m_ of 51 µM, resulting in a catalytic efficiency of 19 592 M^−1^s^−1^ (Figure 5 and Table 2). On the contrary, the D150A variant of SaCntM could use both pyruvate and α-ketoglutarate as substrates. Indeed, although the maximum activity is not reached within the concentration range tested, the kinetic parameters calculated for both pyruvate and α-ketoglutarate are in the same order of magnitude. This single substitution therefore led to a decreased activity when using pyruvate but most of all, significantly increased the activity when using α-ketoglutarate. Even if we take a lower limit for the *k*_cat_ of 1.65 s^−1^ for α-ketoglutarate, this would correspond to a more than 40-fold increased as compared to the WT SaCntM. We then investigated whether the opposite mutation in PaCntM would trigger the same effect on substrate selectivity. Here again, we confirmed that the WT enzyme could only use α-ketoglutarate with a *k*_cat_ of 0.27 s^−1^ and a *K*_m_ of 133 µM. Strikingly, we found that the A153D substitution in PaCntM led to a complete reversal of selectivity, with the variant only being able to use pyruvate and unable to use α-ketoglutarate anymore. This mutation is indeed accompanied by a more than 20-fold increase in *k*_cat_ for pyruvate and ~30-fold decrease for α-ketoglutarate, *i.e.* a complete switch in substrate specificity. Consequently, substrate specificity of CntM towards pyruvate or α-ketoglutarate is mainly governed by this single amino acid (position 150 in SaCntM or 153 in PaCntM): an aspartate ensures the selection of pyruvate whereas an alanine leads to α-ketoglutarate specificity. To our knowledge, this is the first example showing that substrate specificity might be tuned in opine/opaline dehydrogenases family. There are however some examples of redesigned substrate specificity in dehydrogenases such as the production of a highly active malate dehydrogenase starting from lactate dehydrogenase [34]. In this case, the Gln102Arg mutation of *Bacillus stearothermophilus* lactate dehydrogenase led to a shift in *k*_cat_/*K*_m_ with malate so that it equal that of native lactate dehydrogenase for its natural substrate. Examples of site directed mutagenesis studies in the opine/opaline family were centered on the catalytic residues and all led to decreased activities towards substrates and none explored the putative α-ketoacid specificity [12,35].

**Table 2:** Kinetic parameters of SaCntM (WT and D150A mutant) and PaCntM (WT or A153D mutant) activities established for a concentration range of pyruvate and αKG with fixed concentrations of other substrates: 0.2 mM of NADPH and xNA when evaluating SaCntM, and 0.2 mM of NADH and yNA when evaluating PaCntM. The data and the standard errors associated with were generated by SigmaPlot according to the Michaelis-Menten model. (ND: Not Determined). *\**Because the maximum enzyme activity is not reached, these values are not well defined and must be taken with caution.

| **Protein** | **Pyruvate vs αKG** | **$K_m$ (μM)** | **$k_{cat}$ ($s^{-1}$)** | **$k_{cat}/K_m(M^{-1}s^{-1})$** |
| --- | --- | --- | --- | --- |
| SaCntM | pyruvate | $51 \pm 6$ | $1.00 \pm 0.02$ | 19 592 |
| SaCntM | αKG | ND | $0.04 \pm 0.01$ | ND |
| SaCntM:D150A | pyruvate | $681 \pm 115^*$ | $1.23 \pm 0.08^*$ | $1\ 807^*$ |
| SaCntM:D150A | αKG | $2\ 040 \pm 431^*$ | $1.65 \pm 0.22^*$ | $809^*$ |
| PaCntM | pyruvate | ND | $0.02 \pm 0.01$ | N.D. |
| PaCntM | αKG | $133 \pm 39$ | $0.27 \pm 0.02$ | 2 058 |
| PaCntM:A153D | pyruvate | $14\ 986 \pm 2964^*$ | $0.46 \pm 0.07^*$ | 31 |
| PaCntM:A153D | αKG | ND | $0.01 \pm 0.01$ | ND |

### Identification of a novel nicotianamine-like metallophore

Having established the molecular determinant for the α-ketoacid selectivity of CntM, we have proposed simple rules governing the production of nicotianamine-like metallophores: 1-) the presence or absence of a *cntK* homologue leads to the production of D- or L-His respectively (then used by CntL to produce xNA or yNA respectively), and 2-) the presence of an aspartate or an alanine at position 150 (*S. aureus* numbering) results in pyruvate or α-ketoglutarate incorporation, respectively. Applying these rules, we searched for a species capable of producing the missing variant, which would use xNA and α-ketoglutarate, *i.e.* a species possessing a *cntK* homologue and an alanine at position responsible for α-ketoglutarate selectivity. Digging into available genomes *in silico*, we identified *Paenibacillus mucilaginosus* as a good candidate. Indeed, this species carries an A153 in CntM (as *P. aeruginosa*) and possess a *cntK* in its *cnt* operon (as *S. aureus*) (Figure 2B-C).

To check the validity of this hypothesis, genes encoding the CntL and CntM from *P. mucilaginosus* (PmCntL and PmCntM respectively) were amplified from genomic DNA, cloned in expression vectors, purified and used to determine the substrates they consumed *in vitro* (Figure 6). We then assayed enzyme activities using TLC separation and using carboxyl-[^14^C]-labeled SAM. In the TLC assay, [^14^C]-labeled SAM shows a characteristic profile with one strong band and two others bands of much lower intensity (Figure 6 A). The incubation of [^14^C]-labeled SAM with PmCntL led to another prominent band in the presence of D-His, which is not found using L-His. This novel band, migrating below the SAM band, corresponds to the xNA intermediate. This indicates that PmCntL uses D-His and not L-His, and further confirms the link between the presence of a *cntK* gene and the use of D-His by CntL. We then tested the substrate specificity of PmCntM by co-incubating both PmCntL and PmCntM together with diverse substrates: pyruvate or α-ketoglutarate with NADH or NADPH. When using NADPH and D-His with either pyruvate or α-ketoglutarate, we neither detected the SAM nor xNA pattern, but found a novel band migrating just above the SAM in the presence of pyruvate (which corresponds to staphylopine) and just below the SAM in the presence of α-ketoglutarate. Surprisingly, we thus found that PmCntM could use both pyruvate and α-ketoglutarate in this TLC assay. However, TLC experiments were run using a defined incubation time (30 min), which might hinder differences in enzyme efficiency. We therefore determined the enzymatic parameters of PmCntM using the purified protein and a concentration range of pyruvate and α-ketoglutarate with a fixed concentration of others substrates: 0.2 mM of NADPH and 0.2 mM of xNA (Figure 6 B). We found that the catalytic efficiency is 10-fold better for α-ketoglutarate than for pyruvate (10 367 M^−1^s^−1^ and 1 016 M^−1^s^−1^ respectively; Table 3). These data therefore suggest that our hypothesis is valid: CntL and CntM from *P. mucilaginosus* produce an additional variant of opine-type metallophore with D-His, NADPH and α-ketoglutarate. This metallophore has been called bacillopaline as it belongs to the opaline family (Figure 6 C). Moreover, we found that, in addition to using both pyruvate and α-ketoglutarate, CntM from *P. mucilaginosus* is also able to use both xNA and yNA (Figure S9, Table S3). Contrary to human pathogens like *S. aureus*, *P. aeruginosa* or *Y. pestis* in which opine-type metallophores were discovered [5–8], *P. mucilaginosus* is a soil bacteria. The production of opine-type metallophores was shown to be regulated by zinc through the zur (zinc uptake repressor) repressor [6,23], which is likely the case in *P. mucilaginosus*. Therefore, a possible hypothesis is that bacillopaline production could be induced in calcareous soils where zinc bioavailability was shown to be low [37]. In the past decade, *P. mucilaginosus* have been studied for its capacity in wastewater treatment, but also as biofertilizer and plant growth promoting rhizobacteria [38–40]. Indeed, in addition to producing bioflocculants or plant hormones, this species is able to solubilize phosphates, to fix nitrogen and to produce ammonia. Moreover, it is able to produce siderophores, which contributes to plant growth by indirectly preventing the growth of plant pathogens [39,41]. Similarly, we could hypothesize that bacillopaline could contribute in the same way to maintain plant homeostasis.

**Table 3:** Kinetic parameters of PmCntM activities established for a concentration range of pyruvate and αKG with fixed concentrations of others substrates: 0.2 mM of NADPH and 0.2 mM of xNA. The data and the standard errors associated with were generated by SigmaPlot according to the Michaelis-Menten model.

| **Protein** | **Pyruvate vs αKG** | **$K_m$ (μM)** | **$k_{cat}$ ($s^{-1}$)** | **$k_{cat}/K_m$ ($M^{-1}s^{-1}$)** |
| --- | --- | --- | --- | --- |
| PmCntM | pyruvate | $180 \pm 39$ | $0.18 \pm 0.01$ | 1 016 |
| PmCntM | αKG | $42 \pm 5$ | $0.44 \pm 0.01$ | 10 367 |

**Figure 6:**
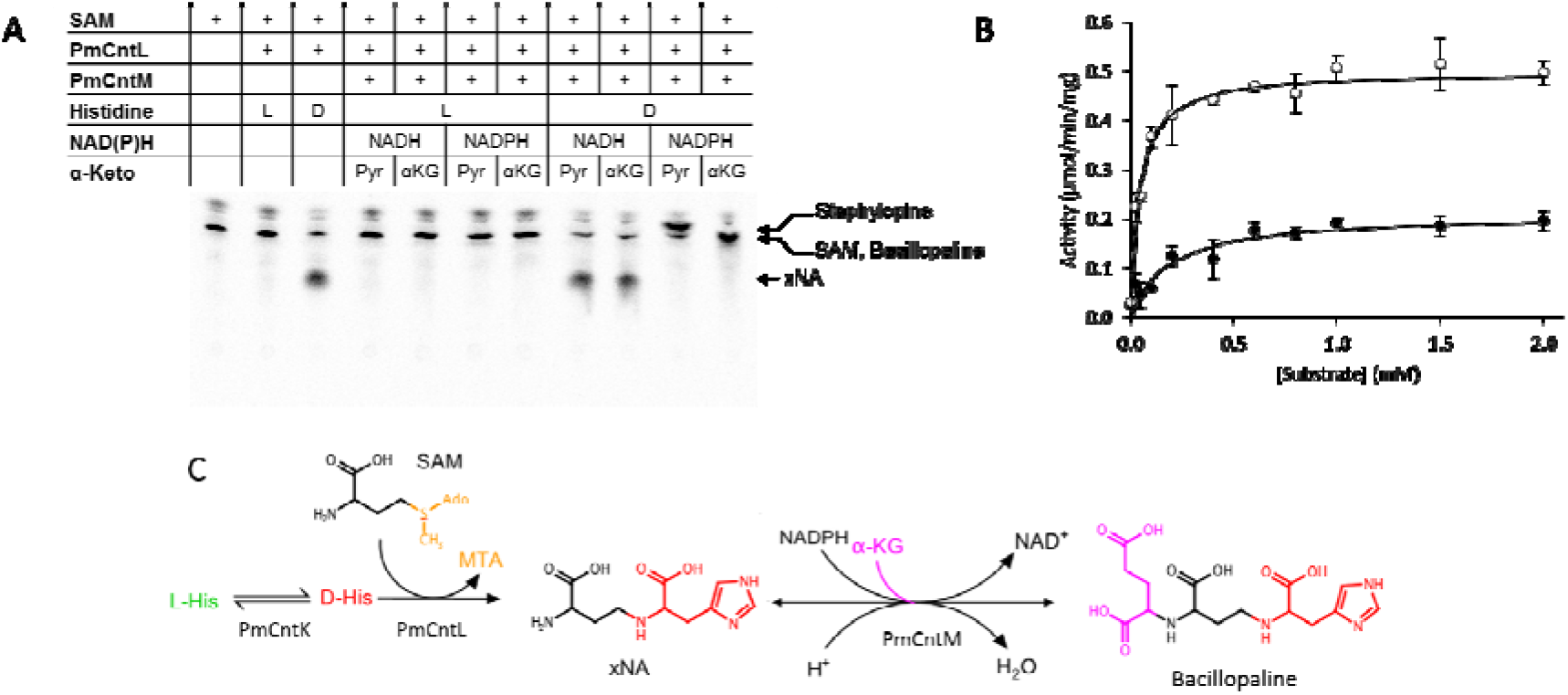
Activity of CntL and CntM from Paenibacillus mucilaginosus. (A) TLC separation of reaction products incubating carboxyl-[^14^C]-labeled with purified enzyme (PmCntL and PmCntM) and various substrates (L-His (L) or D-His (D), pyruvate (Pyr) or α-ketoglutarate (αKG), NADH or NADPH). (B) Activity profile of PmCntM using variable concentrations of pyruvate (black circle) and α-ketoglutarate (white circle) with fixed concentrations of others substrates: 0.2 mM of NADPH and 0.2 mM of xNA. The data points are means of three replicates with standard deviations. The fits are made using the Michaelis-Menten model. (C) Biosynthetic pathway for the assembly of bacillopaline from D-His, SAM and α-ketoglutarate.

In conclusion, studying the substrate specificity of the enzyme catalyzing the last step of the biosynthesis of bacterial nicotianamine-like metallophores allowed us to determine simple rules governing their production. First, the presence or absence of a *cntK* gene leads to the use of respectively D-His or L-His by CntL resulting in the incorporation of respectively xNA or yNA by CntM. Secondly, the presence of an aspartate or an alanine at position 150 on CntM (*S. aureus* numbering) results in pyruvate or α-ketoglutarate incorporation, respectively. Thanks to these simple rules, it is now possible to predict the nature of the nicotianamine-like metallophore produced by all bacteria possessing a *cnt* operon in their genome.

## Acknowledgments

We thank the Agence Nationale de la Recherche (grant ANR-14-CE09-0007-02) for initial support and the association Vaincre la Mucoviscidose (VLM, grant RFI20160501495) for financial support.

## Author contributions

C.L. and P.A. designed the experiments. C.L., C.B., C.H. and L.O. carried out the experiments. C.L., C.B. and P.A. analyzed the data. G.C, R.F synthesized xNA and yNA under the supervision of F.C. C.L. and P.A. wrote the manuscript with contributions from F.C.

