## Supplementary material for "Simple rules govern the diversity of bacterial nicotianamine-like metallophores"

^1^Aix Marseille Univ, CEA, CNRS, BIAM, Saint Paul-Lez-Durance, France F-13108.

^2^Institut des Biomolécules Max Mousseron, IBMM, UMR-5247, CNRS, Université Montpellier, ENSCM , Place Eugène Bataillon, 34095 Montpellier cedex 5, France.

^3^CNRS-UPPA, Laboratoire de Chimie Analytique Bio-inorganique et Environnement, UMR 5254, Hélioparc, 2, Av. Angot 64053 Pau, France.

**SUPPLEMENTARY MATERIALS**

**Chemical synthesis of yNA**

All solvents and reagents for the synthesis were purchased from Sigma Aldrich, Fluka and Alfa Aesar in gradient grade or reagent quality. All reactions involving air-sensitive reagents were performed under nitrogen or argon. Purifications were performed with column chromatography using silica gel (Merck 60, 230–400 mesh). Nuclear magnetic resonance NMR spectra were recorded on a Bruker spectrometer Avance 300 at 600 MHz. Chemicals shifts (*δ*, PPM) are reported from tetramethylsilane with the solvent resonance as internal standard. LC/MS system consisted in a Waters Alliance 2690 HPLC, coupled to a ZQ spectrometer (Manchester, UK) fitted with an electrospray source operated in the positive ionization mode (ESI^+^). All the analyses were carried out using a C18 Chromolith Flash 25 x 4.6 mm column operated at a flow rate of 3 ml/min. A gradient of 0% of 0.1% aqueous TFA (solvent A) to 100% of acetonitrile containing 0.1% TFA (solvent B) was developed over 3 min. Positive-ion electrospray mass spectra were acquired at a solvent flow rate of 100-200 µL/min. Nitrogen was used for both the nebulizing and drying gas. The data were obtained in a scan mode ranging from 200 to 1700 m/z in 0.1 s intervals. A total of 10 scans was summed up to get the final spectrum. Compounds **1** and **2** were purified using a gradient composed of water/acetonitrile with 0.1% TFA at 50 mL/min flow rate performed on a Gilson PLC 2250 preparative apparatus equipped with a C18 Deltapak column (100 mm x 40 mm, 15μm, 100 Å). Purity was determined by RP-Analytic HPLC performed on an Agilent 1220 using a 50 x 4.6 mm Chromolith® High Resolution column. Compounds were separated using a linear gradient system (0 to 100% solvent B in 10 min) using a constant flow rate of 3mL.min^-1^. High-resolution mass spectra (HRMS) were performed by the “Laboratoire de Mesures Physiques” of Montpellier University on a Micromass Q-Tof spectrometer equipped with electrospray source ionization (ESI), using phosphoric acid as internal standard.

**Compound 1**: (*S*)-*tert*-butyl 4-(((*S*)-3-(1*H*-imidazol-5-yl)-1-methoxy-1-oxopropan-2-yl)amino)-2-((*tert*-butoxycarbonyl)amino)butanoate

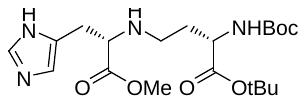

**Compound 2**: (*S*)-2-(((*S*)-4-(*tert*-butoxy)-3-((*tert*-butoxycarbonyl)amino)-4-oxobutyl)amino)-3-(1*H*-imidazol-5-yl)propanoic acid

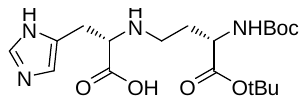

A suspension of H-(L)-His-OMe·2 HCl (305 mg, 1.26 mmol) in MeOH (10 mL) was neutralized with finely powdered NaOH (100.8 mg, 2.52 mmol). After 10 min at room temperature, 300 mg of anhydrous MgSO_4_ were added followed by Boc-(L)-Asa-OtBu (345 mg, 1.26mmol). After stirring the solution for 30 min at room temperature, the reaction was cooled with an ice bath and NaCNBH_4_ (237 mg, 3.78 mmol) was added in portion over 30 min. The reaction mixture was stirred for 3 hours at room temperature then filtered and evaporated under vacuum to remove the solvent. The residue was purified by silica gel chromatography (DCM/MeOH from 98:2 to 96:4) to give the title compound (393 mg, 73%) as a colourless oil.

ESI-MS: 427.0 [M+H]^+^, RP-LC: t_R_ = 0.97 min

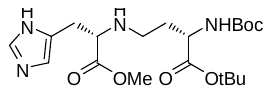

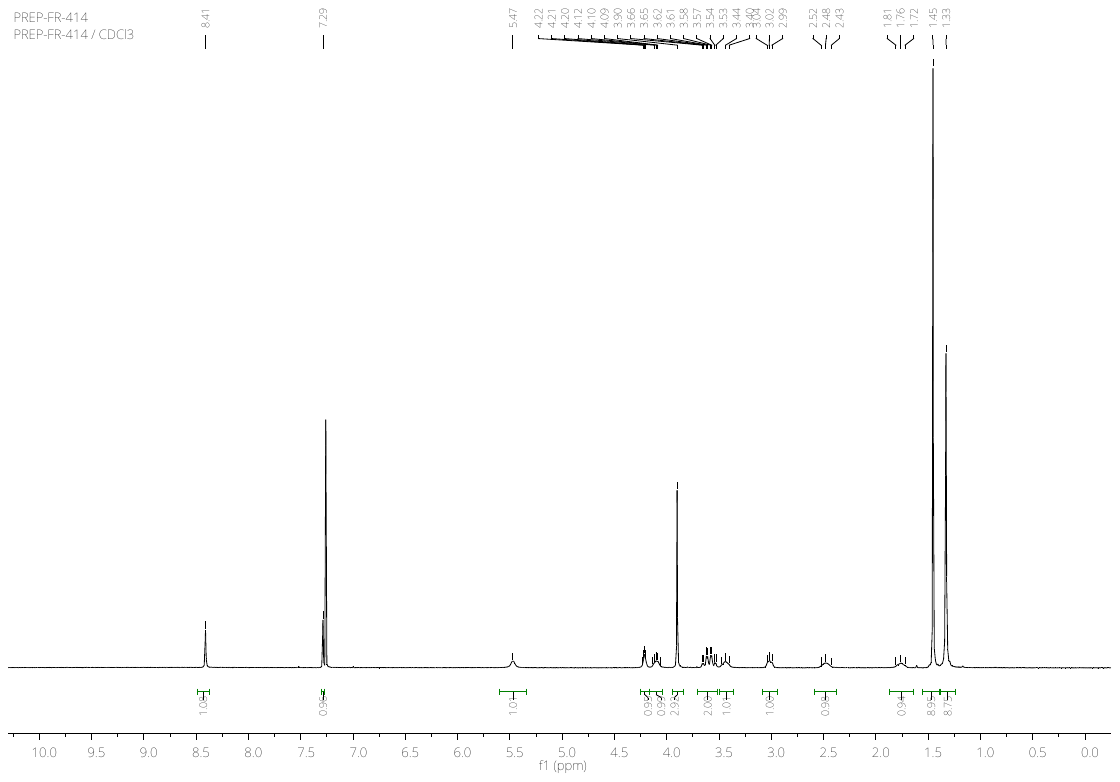

**Figure S1**: 1H NMR spectra of compound **1** (600 MHz, CDCl_3_) δ 8.41 (s, 1H), 7.29 (s, 1H), 5.47 (s br, 1H), 4.23-4.20 (m, 1H), 4.14-4.06 (m, 1H), 3.90 (s, 3H), 3.66-3.54 (m, 2H), 3.53-3.40 (m, 1H), 3.04-2.99 (m, 1H), 2.52-2.43 (m, 1H), 1.81-1.72 (m, 1H), 1.45 (s, 9H), 1.33 (s, 9H).

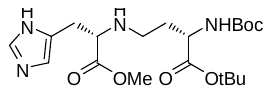

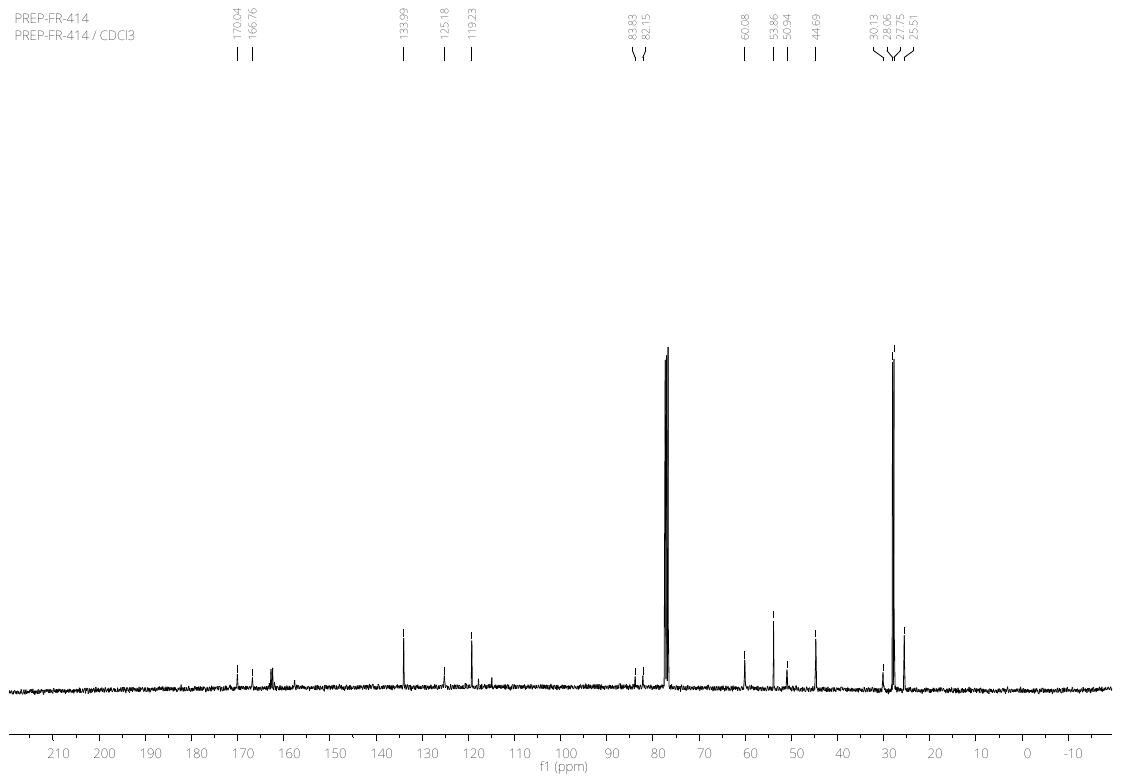

**Figure S2**: ^13^C NMR spectra of compound **1** (150 MHz, CDCl_3_). δ 170.04, 166.76, 133.99, 125.18, 119.23, 83.83, 82.15, 60.08, 53.86, 50.94, 44.69, 30.13, 28.06, 27.75, 25.51.

Compound **1** (155 mg, 0.36 mmol) was dissolved in THF (10 mL) and a solution of LiOH·H_2_O (22.6 mg, 0.54 mmol) in water (3 mL) was added. After 5 h, 0.5 eq. of LiOH·H_2_O were added and the reaction was stirred until completion. The reaction mixture was neutrilized with 1M HCl in dioxane and solvent were removed under vacuum. The residue was purified over preparative HPLC (Buffer A: 0.1% TFA in water; buffer B: 0.1% TFA in acetonitrile, from 0% to 10% of B in 5 min, then to 40% of B in 35 min) and freeze dried to obtain compound **2** as a white powder (91.6 mg, 61%).

ESI-MS: 413.1 [M+H]^+^, RP-LC: t_R_ = 1.09 min

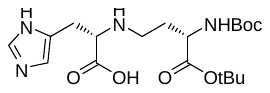

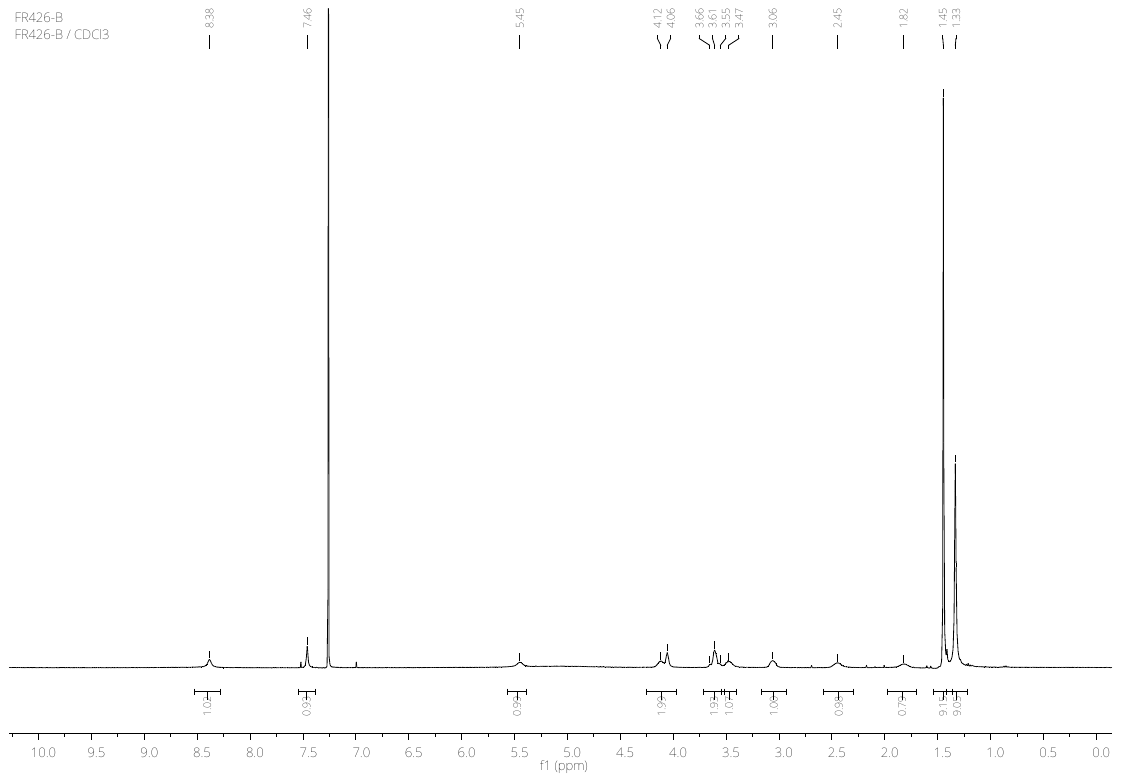

**Figure S3**: 1H NMR spectra of compound **2** (600 MHz, CDCl_3_). δ 8.38 (s, 1H), 7.46 (s, 1H), 5.45 (s br, 1H), 4.12 (br, 1H), 4.06 (br, 1H), 3.6-3.55 (m, 2H), 3.47 (br, 1H), 3.06 (br, 1H), 2.45 (br, 1H), 1.82 (br, 1H), 1.45 (s, 9H), 1.33 (s, 9H).

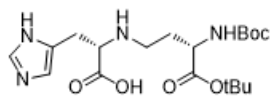

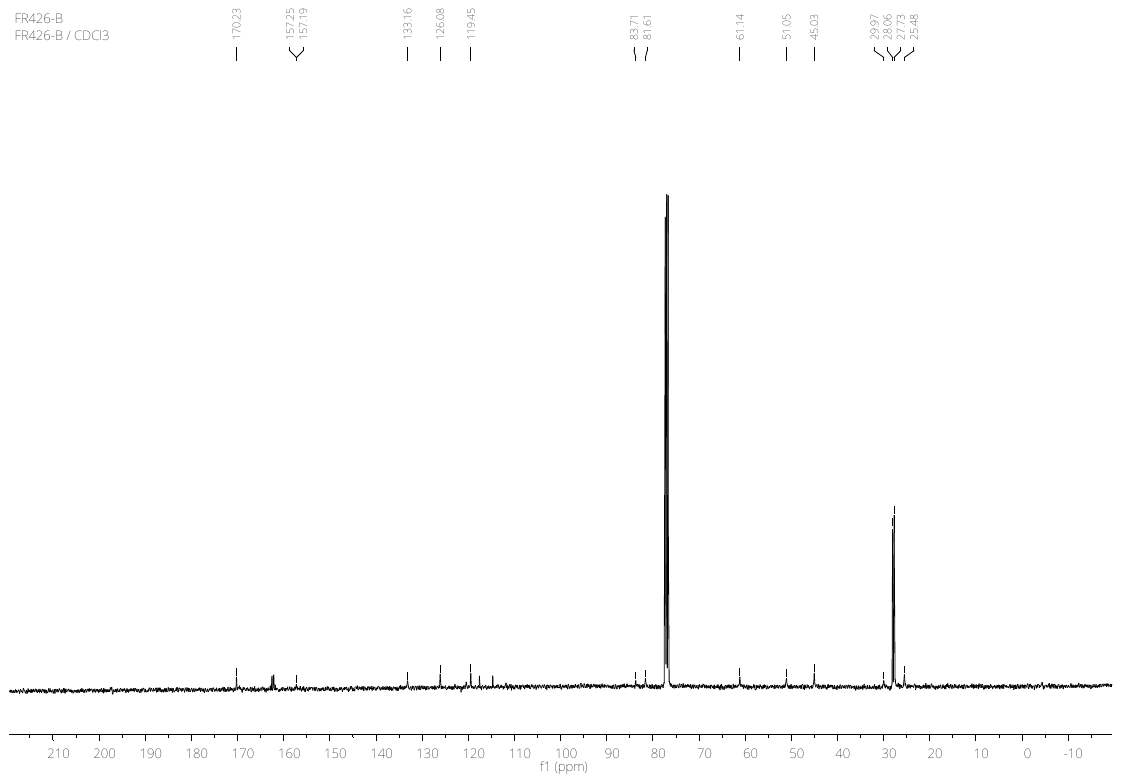

**Figure S4**: ^13^C NMR spectra of compound **2** (150 MHz, CDCl_3_). δ 170.23, 157.25, 157.19, 133.16, 126.08, 119.45, 83.71, 81.61, 61.14, 51.05, 45.03, 29.97, 28.06, 27.73, 25.48.

Compound **2** (55 mg, 0.13 mmol) was dissolved in 1:1 mixture of TFA/DCM (4 mL/mmol) and TIS (3% volume) was added. The mixture was stirred for 3 h at room temperature. Volatiles were removed under reduced pressure. The compounds was dissolved in HCl 0.01 N and freeze dried. This procedure was repeated twice to obtain the hydrochloridric salt **yNA** as a white solid in quantitative yield.

**yNA*3HCl**: 5-((*S*)-2-(((*S*)-3-ammonio-3-carboxypropyl)ammonio)-2-carboxyethyl)-1*H*-imidazol-1-ium chloride

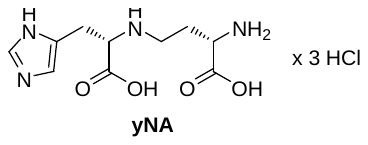

ESI-MS: 257.2 [M+H]^+^, RP-LC: t_R_ = 0.14 min.

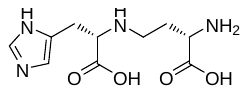

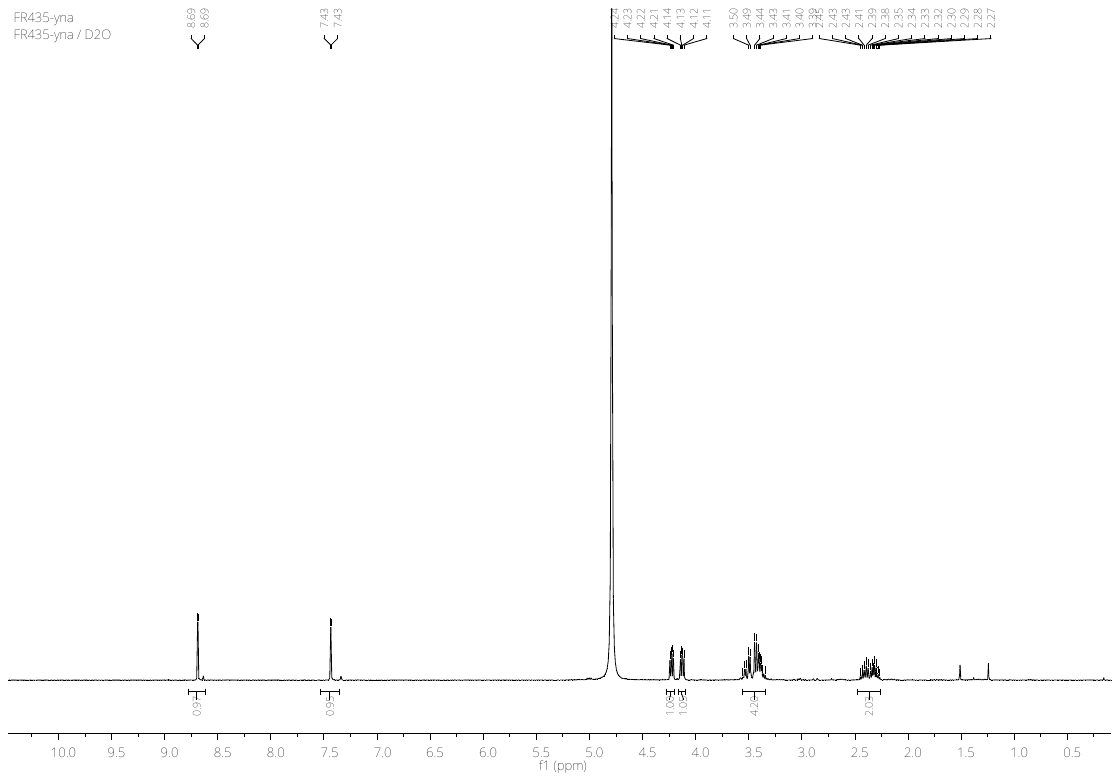

**Figure S5**: 1H NMR spectra of yNA (600 MHz, D_2_O). δ 8.53 (d, *J* = 1.4 Hz, 1H), 7.28 (d, *J* = 1.2 Hz, 1H), 4.07 (dd, *J* = 7.5, 5.3 Hz, 1H), 3.97 (dd, *J* = 8.2, 5.1 Hz, 1H), 3.40-3.16 (m, 4H), 2.35-2.10 (m, 2H).

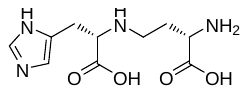

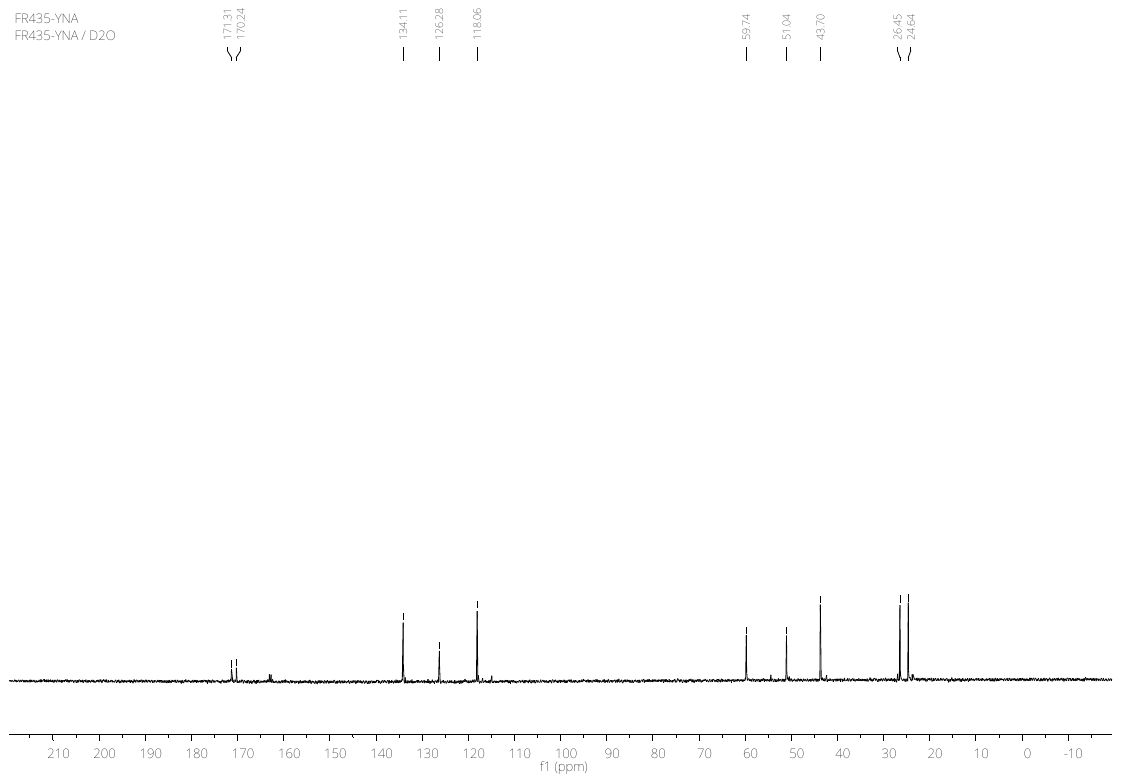

**Figure S6**: ^13^C NMR spectra of xNA (150 MHz, D_2_O). δ 171.31, 170.24, 134.11, 126.28, 118.06, 59.74, 51.04, 43.70, 26.45, 24.64.

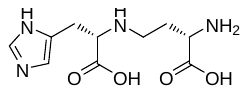

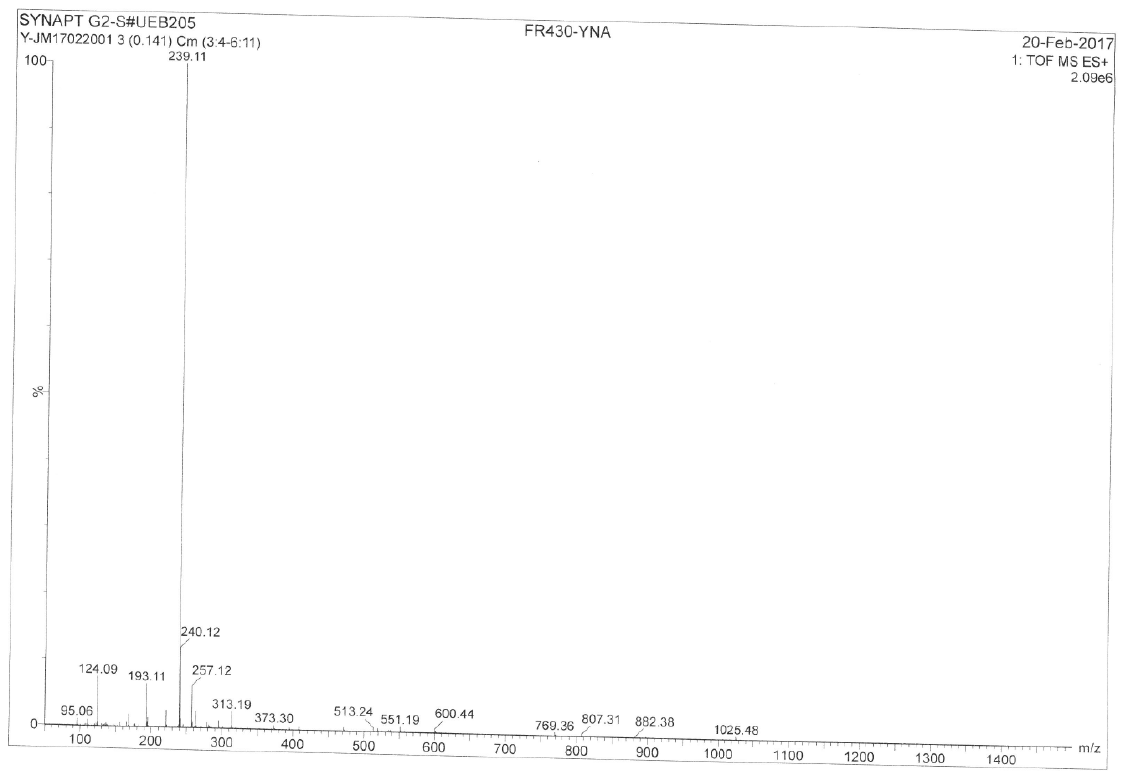

**Figure S7**: HRMS spectrum of yNA (ESI). C_10_H_17_N_4_O_4_ calculated [M+H]: 257.1250, measured: 257.1247.

**SUPPLEMENTARY TABLES AND FIGURES**

| **Name** | **Sequence (5'-3')** |
| --- | --- |
| SaCntM-pET-SUMO Forward | ATGTCTAAATTATTAATGATAGGCACTGGTCCG |
| SaCntM-pET-SUMO Reverse | TTATGAAAGCGTTCTATTGATTTCCAAAAATTTTGTGA |
| SaCntM-pET-101 Forward | CACCATGTCTAAATTATTAATGATAGGCAC |
| SaCntM-pET-101 Reverse | TGAAAGCGTTCTATTGATTTCCA |
| PaCntM-pET-TEV Forward | CATATGAATGCCGCTGATGAGAGC |
| PaCntM-pET-TEV Reverse | CTCGAGTCAGCAGGTCGAGCACCA |
| PmCntL-pET-TEV Forward | GGCATTCCATATGATGAAGACGAAGACACAAGACC |
| PmCntL-pET-TEV Reverse | CCGGAATTCTTAAAGCTGCTCATCATAGGAAC |
| PmCntM-pET-TEV Forward | GGCATTCCATATGATGATGAGCAGCTTTAACCG |
| PmCntM-pET-TEV Reverse | CCGGAATTCTCATGTAAGTTCCCCCGAC |
| SaCntM:R33H Forward | GATATGGTTGGAC***AC***GCCTCAACATC |
| SaCntM:R33H Reverse | GATGTTGAGGC***GT***GTCCAACCATATC |
| SaCntM:D150A Forward | TATCTTGGCG***C***TACACGTATT |
| SaCntM:D150A Reverse | AATACGTGTA***G***CGCCAAGATA |
| PaCntM:A153 Forward | CAGCTACTACGCGG***AC***ACCAAGGTGATCG |
| PaCntM:A153 Reverse | CGATCACCTTGGT***GT***CCGCGTAGTAGCTG |

**Table S1**: Oligonucleotides used in this study.

| **Protein-vector** | **Growth conditions** | **Buffer A** | **Buffer B** | **Imidazole-free buffer B** |
| --- | --- | --- | --- | --- |
| SaCntM-pET-SUMO | 37°C without induction overnight | 20mM Na2HPO4, 300mM NaCl, 15mM Imidazole, pH=8 | 20mM Na2HPO4, 300mM NaCl, 500mM Imidazole, pH=8 | 20mM Hepes, 500mM NaCl, pH=8 |
| SaCntM:D150A-pET-SUMO | 37°C without induction overnight | 20mM Na2HPO4, 300mM NaCl, 15mM Imidazole, pH=8 | 20mM Na2HPO4, 300mM NaCl, 500mM Imidazole, pH=8 | 20mM Hepes, 500mM NaCl, pH=8 |
| SaCntM-pET-101 | 16°C  with induction overnight | 20mM Na2HPO4, 500mM NaCl, 15mM Imidazole, pH=7.5 | 20mM Na2HPO4, 500mM NaCl, 250mM Imidazole, pH=7.5 | 500mM NaCl-Tris, pH=8.5 |
| SaCntM:R33H-pET-101 | 16°C  with induction overnight | 20mM Na2HPO4, 500mM NaCl, 15mM Imidazole, pH=7.5 | 20mM Na2HPO4, 500mM NaCl, 250mM Imidazole, pH=7.5 | 500mM NaCl-Tris, pH=8.5 |
| PaCntM-pET-TEV | 16°C  with induction overnight | 20mM Na2HPO4, 500mM NaCl, 15mM Imidazole, pH=7.5 | 20mM Na2HPO4, 500mM NaCl, 250mM Imidazole, pH=7.5 | 50mM potassium phosphate, 150mM sodium citrate, 20% glycerol, pH=8 |
| PaCntM:A153D-pET-TEV | 16°C  with induction overnight | 20mM Na2HPO4, 500mM NaCl, 15mM Imidazole, pH=7.5 | 20mM Na2HPO4, 500mM NaCl, 250mM Imidazole, pH=7.5 | 50mM potassium phosphate, 150mM sodium citrate, 20% glycerol, pH=8 |
| PmCntM-pET-TEV | 16°C  with induction overnight | 20mM Na2HPO4, 500mM NaCl, 15mM Imidazole, pH=7.5 | 20mM Na2HPO4, 500mM NaCl, 250mM Imidazole, pH=7.5 | 20mM Na2HPO4, 300mM NaCl, pH=8 |
| PmCntL-pET-TEV | 16°C  with induction overnight | 20mM Na2HPO4, 500mM NaCl, 15mM Imidazole, pH=7.5 | 20mM Na2HPO4, 500mM NaCl, 250mM Imidazole, pH=7.5 | 20mM Na2HPO4, 300mM NaCl, pH=8 |

**Table S2**: Growth conditions and buffers used in this study.

Fluorescence (%)

| [NADPH] (µM) | % Moyen WT |
| --- | --- |
| 0 | 14.7019 |
| 1 | 31.0951 |
| 2.5 | 74.3266 |
| 5 | 91.2513 |
| 10 | 94.5304 |
| 50 | 103.178 |
| 100 | 103.324 |
| 200 | 100 |

| [NADPH] (µM) | % Moyen D150A |
| --- | --- |
| 0 | 18.1498 |
| 1 | 27.0228 |
| 2.5 | 60.8788 |
| 5 | 85.9044 |
| 10 | 91.0033 |
| 50 | 104.746 |
| 100 | 104.371 |
| 200 | 100 |

**Figure S8**: Titration of NADPH binding to SaCntM WT (black circle) or to SaCntM:D150A (white circle) followed by fluorescence energy transfer between tryptophan excitation (280 nm) and NADPH emission (450 nm). The signals were measured between 300 and 800nm. We then calculated the mean fluorescence between 400 and 545nm (which correspond to the entire peak of NADPH emission) for a concentration range of NADPH.

PmCntM

Activity (µmol/min/mg)

| [Substrate] (mM) | xNA | yNA | fit xNA non inhib | fit yNA non inhib |
| --- | --- | --- | --- | --- |
| 0 | 0.0851646 | 0.0874415 | 0 | 0 |
| 0.0025 | 0.112643 | 0.156935 | 0.0793103 | 0.151099 |
| 0.005 | 0.135745 | 0.255901 | 0.135294 | 0.237069 |
| 0.0075 | 0.18426 | 0.292752 | 0.176923 | 0.292553 |
| 0.01 | 0.213498 | 0.333477 | 0.209091 | 0.331325 |
| 0.02 | 0.271974 | 0.398262 | 0.2875 | 0.413534 |
| 0.04 | 0.328558 | 0.499466 | 0.353846 | 0.472103 |
| 0.06 | 0.367201 | 0.50869 | 0.383333 | 0.495495 |
| 0.08 | 0.405521 | 0.520296 | 0.4 | 0.508083 |
| 0.1 | 0.437189 | 0.480987 | 0.410714 | 0.515947 |

**Figure S9**: Activity profile of PmCntM using variable concentration of xNA (black circle) and yNA (white circle) with fixed concentrations of others substrates: 0.2 mM of NADPH and 0.2 mM of α-ketoglutarate. The data points are means of three replicates with standard deviations. The fits are made using the Michaelis-Menten model.

| Protein | xNA vs yNA | K_m_ (µM) | k_cat_ (s^-1^) | k_cat_/K_m_ (M^-1^s^-1^) |
| --- | --- | --- | --- | --- |
| PmCntM | xNA | 12 ± 1 | 0.40 ± 0.02 | 33 383 |
| PmCntM | yNA | 7 ± 1 | 0.48 ± 0.01 | 72 572 |

**Table S3** : Kinetic parameters of PmCntM activities established for a concentration range of xNA and yNA with fixed concentrations of others substrates: 0.2 mM of NADPH and 0.2 mM of α-ketoglutarate. The data and the standard errors associated with were generated by SigmaPlot according to the Michaelis-Menten model.
